# Two species and two karyotypes? Cytogenetic differentiation between *Passiflora foetida* L. and *P. vesicaria* L. (Passifloraceae)

**DOI:** 10.1101/2024.10.23.619917

**Authors:** Bruna Zirpoli, Arthur Roberto Monteiro da Silva, Pablo E. Rodriguez, Bruno Leal Viana, Jefferson Rodrigues Marciel, Andrea Pedrosa-Harand, Mariela A. Sader

## Abstract

The genus *Passiflora* L. (Passifloraceae) is widely distributed, primarily in the Neotropics, and comprises over 500 species divided into four main subgenera. The subgenus *Passiflora* is the most economically relevant and has a basic chromosome number of *x* = 9, with *P. foetida* L. being the sole species in the subgenus having a chromosome number of *n* = 10. However, *P. foetida* exhibits substantial morphological variation among its varieties, and recently, two of these varieties were reclassified as *Passiflora vesicaria* L. and *P. vesicaria* var. *galapagensis*. This study aimed to cytogenetically characterise both species by estimating genome sizes using flow cytometry, mapping heterochromatic regions by CMA/DAPI staining, and identifying 5S and 35S ribosomal DNA sites using fluorescent *in situ* hybridization (FISH). The genome sizes were similar across the three accessions analyzed, varying from 2C = 1.00 ± 0.03 pg for *P. vesicaria* var. *galapagensis* to 2C = 1.16 ± 0.01 pg for *P. vesicaria*, indicating small genomes. All three accessions showed 2*n* = 20, confirming the count for *P. foetida* and revealing the chromosome numbers of the two *P. vesicaria* accessions for the first time. Six CMA^+^ bands were observed in *P. foetida*, four in *P. vesicaria* var. *galapagensis*, and two in *P. vesicaria*, all co-localised with the 35S rDNA sites. FISH with 5S and 35S rDNA showed six pericentromeric 35S sites of different intensities in *P. foetida* and four interstitial 5S sites, with one individual having only two interstitial 5S sites, demonstrating a polymorphism at the intrapopulation level not yet recorded for the genus or the species. *Passiflora vesicaria* exhibited four 35S sites also pericentromeric, as well as two 5S sites, while *P. vesicaria* var. *galapagensis* presented four pericentromeric 35S sites and two 5S sites. Therefore, despite these species having the same chromosome number (*n* = 10) and similar genome sizes, there are karyotypic variations in the number of CMA^+^ bands and rDNA sites, allowing their discrimination.

## Introduction

The *Passiflora* L. genus is the largest in the family Passifloraceae, consisting of around 560 species, mostly climbers, lianas, and woody plants, although it also includes species with shrubby or tree-like habits (Koch & Ilkiu-Borges, 2016; Krosnick et al., 2013). The genus is distributed mainly in the Neotropics (Ulmer & MacDougal, 2004) and has coevolutionary relationships with many organisms, including pollinators (Ocampo Perez & Coppens d’Eeckenbrugge, 2017). Due to its wide morphological variation, the genus was divided into 22 subgenera (Killip, 1938); however, more recently, they have been grouped into four main subgenera: *Astrophea* (DC.) Mast., *Decaloba* (DC.) Rchb., *Deidamioides* (Harms) Killip, and *Passiflora* Feuillet & MacDougal (Ulmer and MacDougal, 2004).

The *Passiflora* subgenus comprises about 250 species, making it the largest subgenus of *Passiflora* (Cervi & Imig, 2013). It includes climbers and lianas with relatively large flowers, with uni-to multiseriate coronal filaments, five stamens, and a stigma and ovary raised on a column called the androgynophore. It is widely distributed, mainly in the tropics, and is the most well-known subgenus due to its economic importance (Ulmer & MacDougal, 2004). *Passiflora foetida*, known as stinking passionflower, wild maracuja, bush passion fruit, wild water lemon, stoneflower, love-in-a-mist, or running pop, was the first lineage to diverge within the *Passiflora* subgenus and is considered the sister species of all other species in the subgenus (Cauz-Santos et al., 2020). It is a widely distributed species (Patil et al., 2015), with high rates of reproduction and dispersion, and is therefore considered invasive in some countries (Cowie & Werner, 1993; Hopley et al., 2021). Some studies suggest that *P. foetida* has the potential to be used for gene introgression in commercial plants (Vijay et al., 2021) and has potential antifungal properties (Elangovan et al., 2022), in addition to the pharmacological uses known for the entire genus (Chiavaroli et al., 2020; Leal et al., 2022).

The species exhibits a wide morphological diversity, and it is possible to distinguish six varieties. Furthermore, taxa that were previously considered as varieties of *P. foetida* were elevated to the species level based solely on morphological data, namely *P. vesicaria* and *P. vesicaria* var. *galapagensis* (Vanderplanck, 2015). In a characterization of *Passiflora foetida* aimed at investigating the origin of invasive accessions in Australia, three lineages were recognised: Brazil (Clade I), Caribbean (Clade II), and Ecuador (also Clade II) (Hopley et al., 2021). However, it was not possible to differentiate between their varieties, and all taxa were treated as *P. foetida lato sensu*. In addition to morphological and molecular characterizations, karyotypic characterization can aid in the delimitation of these taxa using a cytotaxonomic approach (Guerra, 2008).

Among the main cytogenetic characteristics used in cytataxonomy are chromosome number, genome size, heterochromatic bands, and number and location of 35S and 5S ribosomal DNA (rDNA) sites (Guerra, 2012). For the identification of heterochromatic bands, the CMA/DAPI double staining is commonly used in plants, which stains respectively the regions rich in GC and AT (Cordeiro et al., 2022). The 5S and 35S rDNA are tandemly repeated sequences that encode the 5S, 5.8S, 18S, and 25-28S rRNAs, of extreme importance for ribosome assembly (Richard et al., 2008) and are used as cytomolecular markers in studies of chromosomal evolution due to its conserved sequences (Jiang et al., 2019).

Although *Passiflora* presents a variable chromosome number among its subgenera (Sader et al., 2019a), most of the subgenus *Passiflora* has meta- and submetacentric chromosomes with a basic chromosome number *x* = 9 (Sader et al., 2019b), and rDNA sites at terminal or subtelomeric positions (Coelho et al., 2016; Dias et al., 2020). However, the analyzed accessions of *P. foetida* showed *n* = 10 (Barros et al., 2021; Ferreira et al., 2020; Melo et al., 2001; Mikovski et al., 2021), and presented 35S rDNA sites at proximal positions (Melo & Guerra, 2003). Nevertheless, none of the studies were the materials classified at variety level and intraspecific variability was not considered.

Therefore, the present study aimed to characterise the karyotypes of four accessions of the species *P. foetida* and P. *vesicaria*, a recently separated species from *P. foetida*, using CMA/DAPI double staining, mapping the ribosomal DNA 5S and 35S, and estimating genome sizes to enable taxonomic and evolutionary discussions regarding the *Passiflora* subgenus.

## Materials and Methods

### Plant Material

Seeds from two accessions of *P. vesicaria* and one of *P. foetida* species were obtained from the Recife Botanical Garden. A fourth accession, from *P. foetida*, was obtained from the *Passiflora* Germplasm Bank “Flor da Paixão” located at Embrapa Cerrados in Planaltina, DF. The plants were grown in the Experimental Garden of the Laboratory of Plant Cytogenetics and Evolution (UFPE) and multiplied by cuttings. Vouchers were deposited in the UFP Herbarium - Geraldo Mariz (Table 1). The specimens are illustrated in Fig. 1.

**Figure 1.**
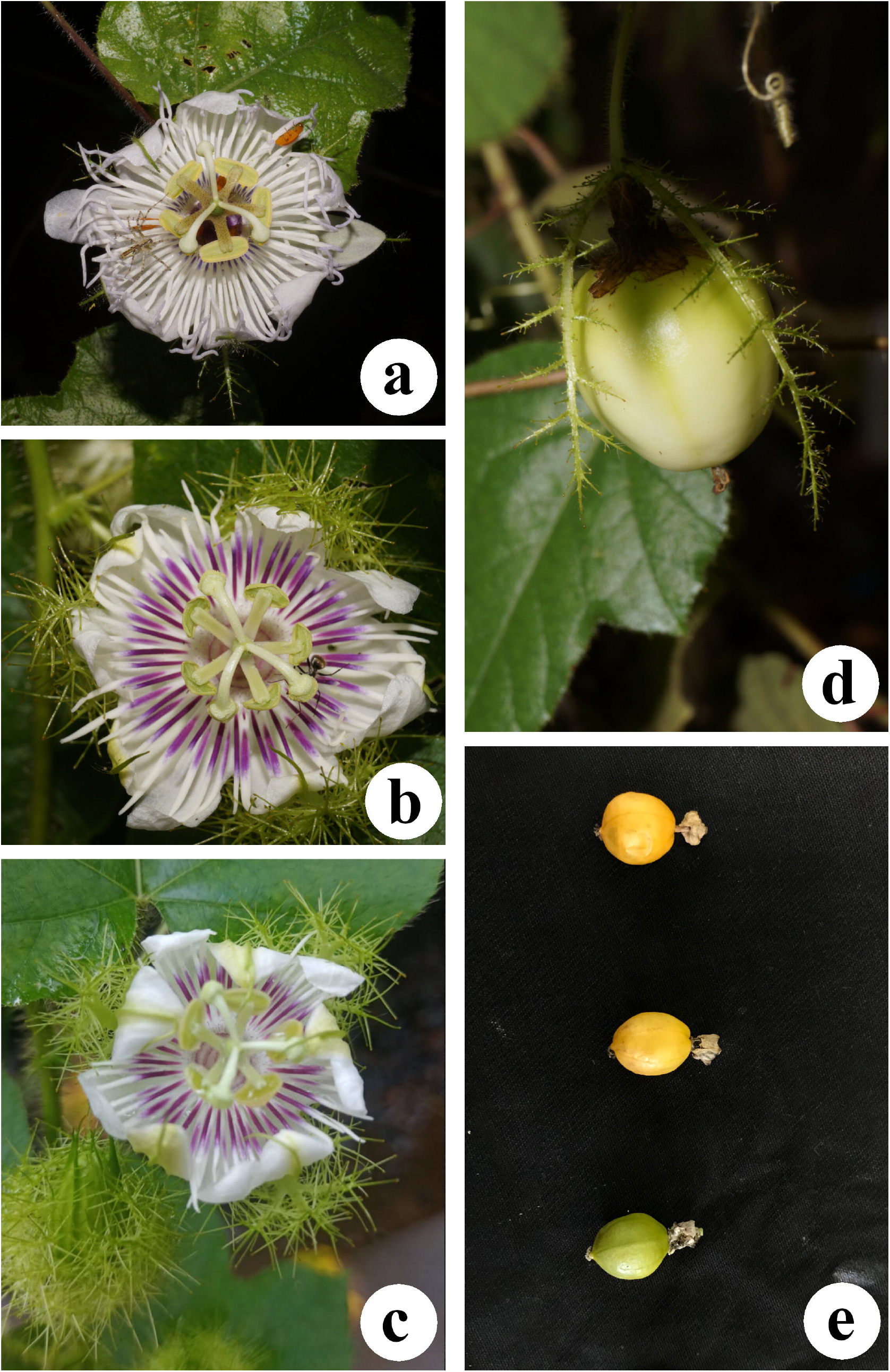
Morphological plate of *Passiflora foetida* and *P. vesicaria* accessions analysed. a-c. Flowers: a. *P. foetida*; b. *P. vesicaria* var. *vesicaria*; c. *P. vesicaria* var. *galapagensis*; d-e. Fruits: d. *P. foetida*; e. *P. vesicaria* var. *galapagensis*.

### Cytogenetic Analysis

Seeds of the species were germinated on moist filter paper in Petri dishes after mechanical scarification and incubation for 30 min in Promalin® (12gr/lt) (Sumitomo Chemical Brasil) at room temperature. Root tips from germinated seeds or cuttings were pre-treated with 8-hydroxyquinoline (8-HQ) for 4.5 hours at 10°C, fixed in Carnoy 3:1 (ethanol: glacial acetic acid) for at least one hour at room temperature, and stored in a freezer at -20°C for later use. For mitotic preparations, the roots were washed twice with distilled water and digested in a solution containing 2% cellulase (Onozuka) and 20% pectinase (Sigma) (v/v) at 37°C for one to three hours.

Chomosome preparations were performed according to the air-drying protocol with modifications described by Ribeiro et al. (2017). Chromosome banding by double staining with the fluorochromes Chromomycin A3 (CMA) and 4’6-diamidino-2-phenylindole (DAPI) was carried out according to Vaio et al. (2018). Fluorescent *in situ* hybridizations were performed according to Fonsêca et al., (2010) with pre-labeled (Cy3) oligonucleotides for 5S rDNA (5S-PLOPs, Waminal et al., 2018). The wheat clone p*Ta*71 (25-28S, 5.8S, and 18S rDNA; Gerlach and Bedbrook, 1979) labeled by nick translation with Alexa-488-dUTP was used as probe for the 35S rDNA sites. Images were captured using a Leica DM5500B epifluorescence microscope with Leica LAS AF capture system. Adobe Photoshop CS6 was used for uniform adjustments of the images for brightness and contrast.

### Flow cytometry

For the cytometry analyses, fresh leaves from two individuals of each accession under cultivation were used in triplicate for genome size estimations simultaneously with the internal standard (*Lycopersicon esculentum* Mill. 2C = 1.96 pg, Doležel et al. 1998), according to the protocol revised by Pellicer & Leitch (2014), in 1 mL of Marie Buffer. A 30 μm filter was used, and 50 mg/mL of propidium iodide was added as a DNA intercalating fluorochrome. A CyFlow Space (Sysmex, Norderstedt, Germany) flow cytometer was used for data acquisition, and relative fluorescence histograms were generated with Flomax v.2.3.0. software (Sysmex, Norderstedt, Germany).

## Results

The four accessions of the two analyzed species presented the same chromosome number, 2*n* = 20, with metacentric and submetacentric chromosomes (Table 1, Fig. 2). These counts corroborated previous records of *P. foetida* (Guerra, 2001; Melo & Guerra, 2003) and added new chromosomal counts for *P. vesicaria* and *P. vesicaria* var. *galapagensis*.

**Figure 2.**
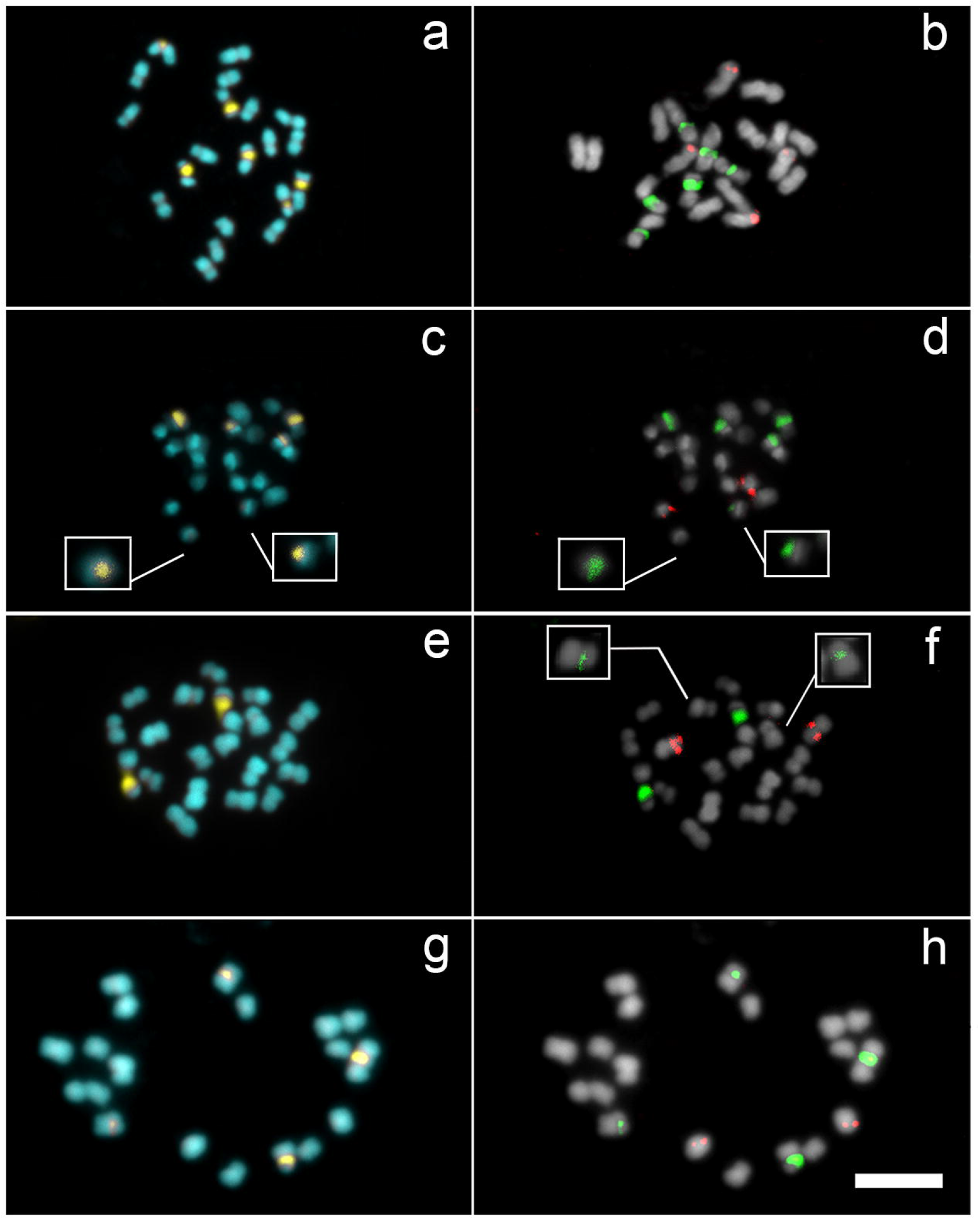
Distribution of CMA (yellow) and DAPI (blue) heterochromatic bands and 5S (Cy3, in red) and 35S (Alexa, in green) rDNA sites (b, d, f, and h) in *Passiflora foetida* and *P. vesicaria*. (a-d) *P. foetida*; (e-f) *P. vesicaria* with insets of the two small 35S sites; (g-h) *P. vesicaria* var. *galapagensis*. Scale bar in corresponds to 10 μm.

The CMA/DAPI double staining revealed CMA-positive heterochromatic bands in three pairs of chromosomes in the two accessions of *P. foetida*, totaling six bands in the pericentromeric region (Fig. 2a), while *P. vesicaria* showed only one strong CMA^+^ band in the pericentromere (Fig. 2e). *P. vesicaria* var. *galapagensis* showed one strong and one weaker pair of CMA^+^ bands (Fig. 2g), resembling the *P. vesicaria* karyotype (Table 1).

FISH with ribosomal DNAs, *P. foetida* showed two pairs of 5S rDNA sites in interstitial positions and three pairs of proximal 35S rDNA sites (Table 1, Fig. 2b, 3a), corresponding to the CMA^+^ bands. However, the accession obtained from the *Passiflora* Germplasm Bank “Flor da Paixão’’ showed only one pair of 5S rDNA (Fig. 2d), revealing an absence of 5S rDNA in the third pair of chromosomes (Fig. 3b). *Passiflora vesicaria* had one pair of interstitial 5S rDNA and two pairs of 35S rDNA, one strong and the other significantly weaker (Fig. 2f, 3c). The stronger 35S rDNA signals were co-located with the CMA^+^ bands, but the weaker pair was not visible as CMA^+^ bands (Fig. 4). *Passiflora vesicaria* var. *galapagensis* presented, as expected, four 35S rDNA sites (Fig. 2h, 3d) given the CMA/DAPI pattern observed previously (Fig. 2g). There was no difference in the number of rDNA sites compared to the typical *P. vesicaria* accession, but only in the size of the 35S rDNA sites (Table 1, Fig. 2f and h).

**Figure 3.**
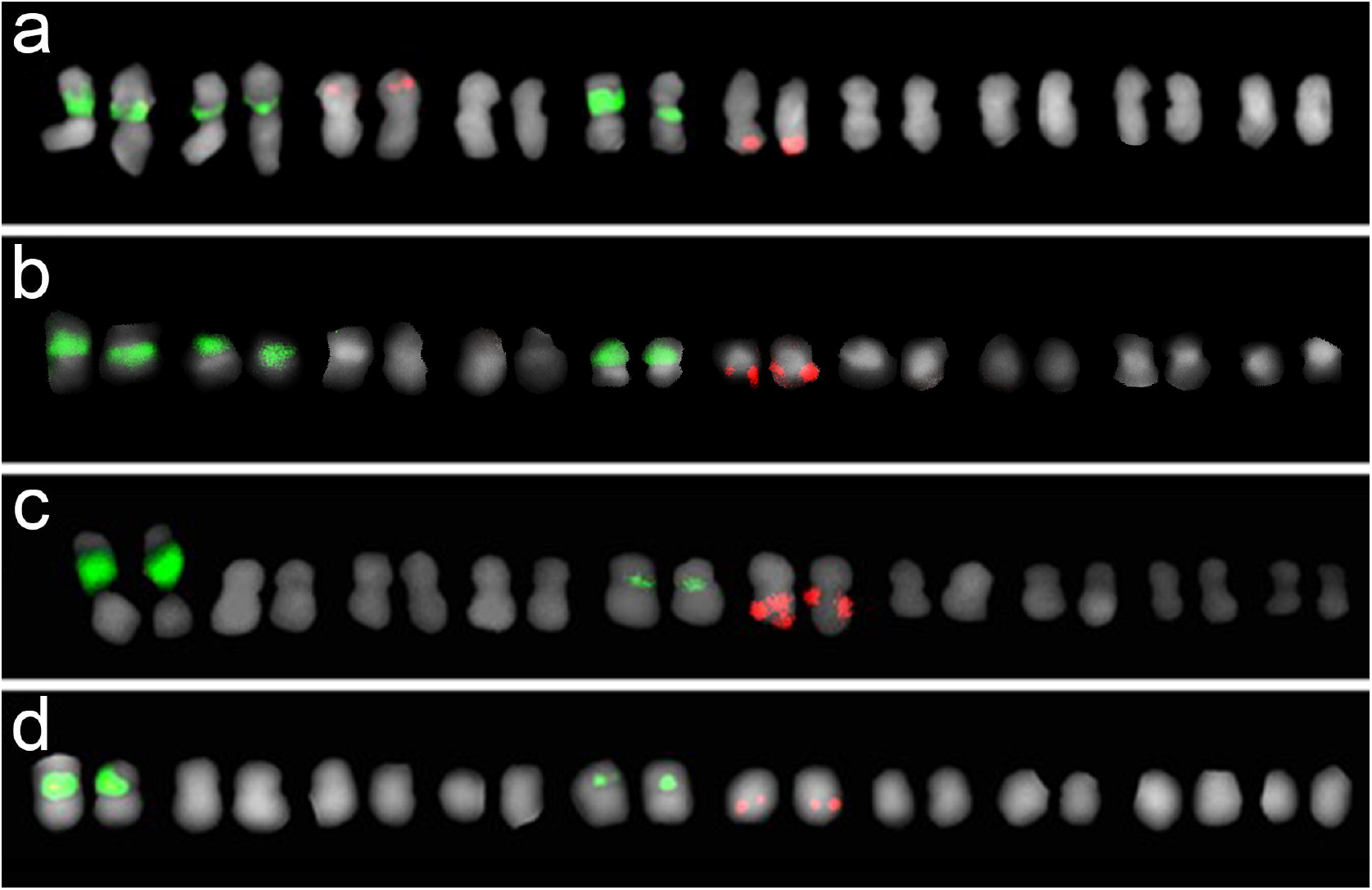
Karyograms of *Passiflora foetida* (a, b), *P. vesicaria* (c), and *P. vesicaria* var. *galapagensis* (d) with rDNA 5S (red) and 35S (green) sites.

**Figure 4.**
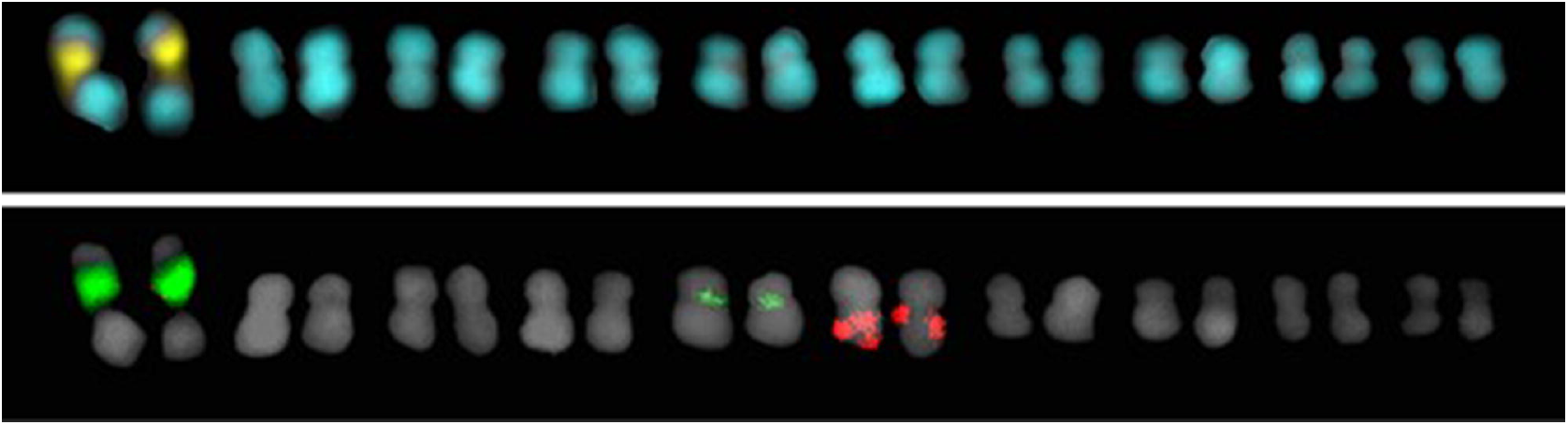
Comparative karyograms of CMA/DAPI double staining (yellow and blue) and 5S (red) and 35S (green) rDNA FISH of a *Passiflora vesicaria* cell, showing colocalization of the CMA^+^ bands (yellow) and the strongest 35S rDNA sites (green).

The genome size estimates indicated 2C = 1.07 ± 0.07 pg for *P. foetida*, 2C = 1.16 ± 0.01 pg for *P. vesicaria*, and 2C = 1.00 ± 0.03 pg for *P. vesicaria* var. *galapagensis*. The three accessions showed a small variation in genome size (Table 1, Fig. 5).

**Figure 5.**
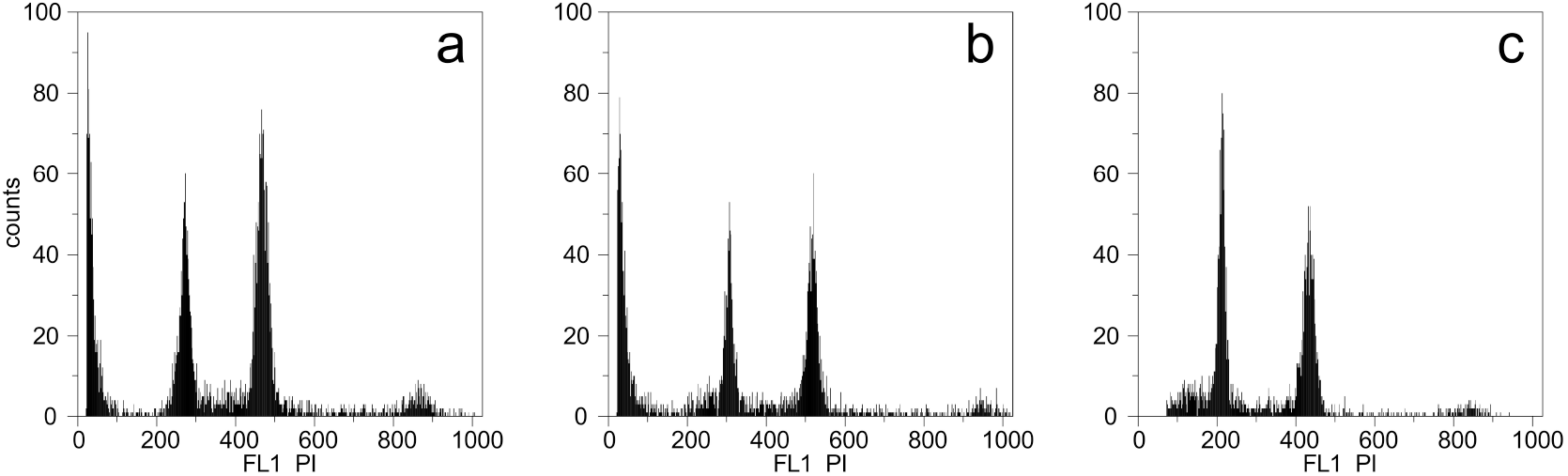
Flow cytometry histograms. The first peak in each histogram corresponds to the species (a) *P. foetida*, (b) *P. vesicaria*, and (c) *P. vesicaria* var. *galapagensis*, while the second peak corresponds to the control, *Lycopersicon esculentum*.

## Discussion

In this study, we presented the cytogenetic characterization of two varieties of *P. vesicaria* and compared them with similar data of two accessions of *P. foetida*, corroborating chromosome counts for *P. foetida* (Barros et al., 2021; Ferreira et al., 2020; Melo & Guerra, 2003; Melo et al., 2001; Mikovski et al., 2021) and providing new counts for *P. vesicaria*. The variations in chromosome number previously reported for *P. foetida*, 2*n* = 18 (Ammal, 1945) and 2*n* = 22 (Bowden, 1945; Harvey, 1966), were probably due to extended, proximal secondary constrictions or uncertain counts.

The genome size in *Passiflora* varies considerably among its species, ranging from 2C = 0.21 pg in *P. organensis* to 5.36 pg in *P. quadrangularis*, with larger genomes in the subgenus *Passiflora* (Souza et al., 2004; Yotoko et al., 2011). The values found in this study do not suggest significant variation between the two species and are similar to those previously described for *P. foetida* (2C = 0.96 pg) (Yotoko et al., 2011). Their relatively small genome is compatible with the observed genome size increase in other species of the subgenus *Passiflora* (Sader et al., 2019b).

Although all four taxa presented similar genome sizes and 2*n* = 20, corroborating their close relationship (Vanderplank, 2013), differences in the number of CMA^+^ heterochromatic bands and 35S and 5S rDNA sites allowed the cytogenetic discrimination of the species, supporting the classification of *P. vesicaria* as a separate species. As for the variety *P. vesicaria* var. *galapagensis*, the data showed similarities with *P. vesicaria*, with a slight difference in the intensity of a 35S rDNA site that reflected in a less visible CMA^+^ band in the typical variety, which has also been observed in *Citrus reticulata* cv. Cravo (Da Costa Silva et al., 2015).

The variation in 5S rDNA recorded in one of the accessions of *P. foetida* (UFP90688) results from an intraspecific polymorphism that had not been reported for the subgenus *Passiflora*. Chromosomal polymorphism may result either from “silent” structural changes without phenotypic expression (Takahashi et al., 2018) or contribute to its genetic variability, being neutral or related to its better adaptation to the environment (Tabatabeaei et al., 2019). This accession was identified as *P. foetida* var. *foetida* by morphological analysis, although no phenotypic differences were observed between both *P. foetida* accessions. Regarding its morphology, *P. foetida* has colourful flowers (Fig. 1), ranging from purple to white allowing the identification of six varieties.

Morphological traits, such as flower colour, fruit and seed shape, and the number of leaf lobes (Fig. 1b-c), have supported the categorization of *P. vesicaria* varieties into the species status (*P. vesicaria* and *P. vesicaria* var. *galapagensis*; Vanderplanck, 2013), even though they do not present significant morphological differences besides *P. vesicaria* var. *galapagensis* is geographically very isolated. They also do not show significant differences in their fruits when compared among themselves and to *P. foetida* (Fig. 1d-e).

The variations in the amount of rDNA sites are taxonomically significant within the subgenus; other more distantly related species showed different numbers of sites, such as *P. edulis* (four 35S rDNA sites) (Melo & Guerra, 2003; Sader et al., 2019a) and *P. watsoniana* (six 35S rDNA sites) (Dias et al., 2020), making it possible to separate them based on this trait (Melo & Guerra, 2003). The position of rDNA sites may also vary among species, although it has been shown to be conserved amongst closer species. While in *P. edulis*, as in many other species of the *Passiflora* subgenus, the 35S rDNA sites were observed in terminal position (Coelho et al., 2016; Dias et al., 2020), in *P. foetida* and *P. vesicaria* they were proximal, as also present in *P. misera* and *P. tricuspis* of the *Decaloba* subgenus (Melo & Guerra, 2003).

Considering the basal phylogenetic position of *P. foetida* in the subgenus (Cauz-Santos et al., 2020) and a likely ancestral karyotype for the genus with *x* = 12 (Melo & Guerra, 2021), the observed *x* = 10 of *P. foetida* and *P. vesicaria* probably represents an intermediate stage resulting from descending dysploidy, which later originated the most frequent haploid number in other species of the subgenus, *x* = 9. Thus, *x* = 10 should be considered the ancestral basic chromosome number for the *Passiflora* subgenus (Fig. 6). In this scenario, the proximal sites could be the result of the fusion of chromosomes with terminal sites, as has been observed in species of the subgenus *Astrophea*, with *n* = 12 (Melo & Guerra, 2003). However, most species in the subgenus have *n* = 9 and 35S rDNA terminal sites (Dias et al., 2020), suggesting more complex structural rearrangements in the karyotypes during dysploidy events. In the *Senna* Mill. group, when comparing *Senna occidentalis* (L.) Link and *S. tora* (L.) Roxb, descending dyploidy occurred through various structural rearrangements, involving whole-genome duplications and chromosomal fusions (Waminal et al., 2021). The alternative hypothesis for this change in the location of rDNA sites would be rDNA transposition independent of structural rearrangements, as reported in *Triticeae* (Dubcovsky and Dvorak, 1995; Raskina et al., 2004a, b). A broader sampling of various individuals from different locations of each variety of both species would be necessary to analyse the extent and possible causes of the rDNA variability and if this trait could also differentiate the taxa at the intraspecific level.

**Figure 6.**
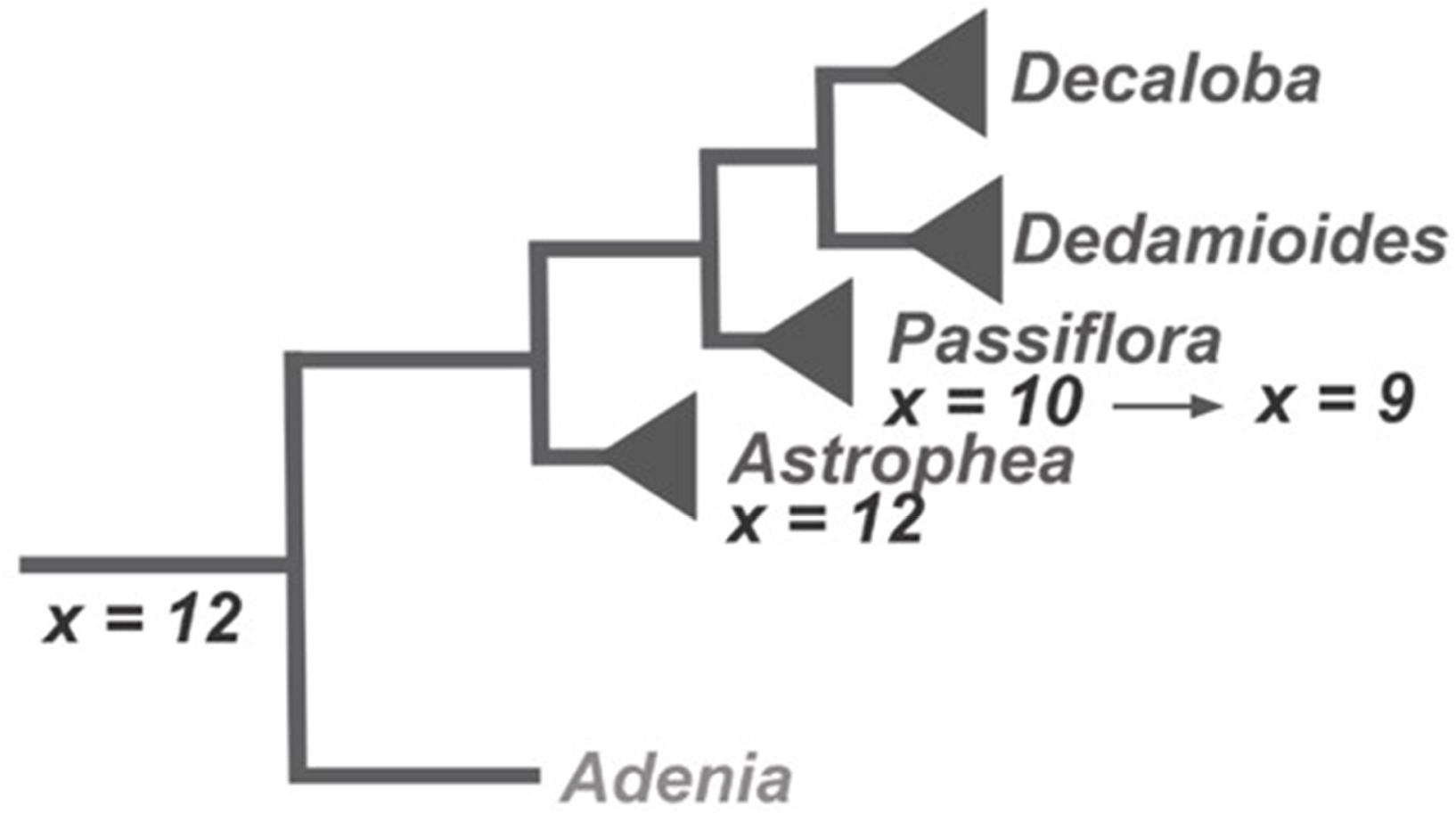
Simplified representation of the phylogenetic relationships and reconstruction of the ancestral chromosome number and its changes in subgenus *Passiflora*, initially from *x* = 12 to *x* = 10 and subsequently to *x* = 9, the most common number currently found in the subgenus. Modified from Cauz-Santos et al. (2020) and Melo & Guerra (2021).

## Acknowledgements

We would like to thank Nilton Tadeu V. Junqueira from the Passiflora Germplasm Bank “Flor da Paixão” located at Embrapa Cerrados in Planaltina, DF, and Marcelo Guerra from UFPE, Brazil, for providing plant material for this study. We are grateful to the Fundação de Amparo à Ciência e Tecnologia de Pernambuco (FACEPE); the Conselho Nacional de Desenvolvimento Científico e Tecnológico (CNPq), Brazil; Consejo Nacional de Investigaciones Científicas y Técnicas (CONICET); and Agencia Nacional de Promoción Científica y Tecnológica (FONCyT), Argentina, for financial support.

## Author Contributions

BZ performed experiments, data analysis and wrote the first version of the manuscript. ARMS performed FISH experiments and assisted with writing the manuscript, PR performed flow cytometry analysis and assisted with writing the manuscript; BLV and JRM provided plant material, morphological characterization and taxonomic identification. APH and MS designed the experiments, analysed data and assisted with writing the manuscript. All authors have read and agreed to the final version of the manuscript.

