## Supplementary material for "Two species and two karyotypes? Cytogenetic differentiation between *Passiflora foetida* L. and *P. vesicaria* L. (Passifloraceae)": Table 1

| Species | Voucher | 2*n* | 2C (pg)/ cv | CMA+ bands | 5S rDNA | 35S rDNA |
| --- | --- | --- | --- | --- | --- | --- |
| *P. foetida* var*. foetida* | UFP85988 | 20 | 1,07 ± 0,07/ 3,32 | 6p | 4i | 6p |
| *P. foetida* var*. foetida* | UFP90688 | 20 | - | 6p | 2i | 6p |
| *P. vesicaria* | UFP89157 | 20 | 1,16 ± 0,01/ 3,13 | 2p | 2i | 4p |
| *P. vesicaria* var. *galapagensis* | UFP85083 | 20 | 1,00 ± 0,03/ 4,71 | 4p | 2i | 4p |
